# High temperatures drive offspring mortality in a cooperatively breeding bird

**DOI:** 10.1101/2020.05.31.126862

**Authors:** Amanda R. Bourne, Susan J. Cunningham, Claire N. Spottiswoode, Amanda R. Ridley

## Abstract

An improved understanding of life history responses to current environmental variability is required to predict species-specific responses to anthopogenic climate change. Previous research has suggested that cooperation in social groups may buffer individuals against some of the negative effects of unpredictable climates. We use a 15-year dataset on a cooperative-breeding arid-zone bird, the southern pied babbler *Turdoides bicolor*, to test i) whether environmental conditions and group size correlate with survival of young during three development stages (egg, nestling, fledgling), and ii) whether group size mitigates the impacts of adverse environmental conditions on reproductive success. Exposure to high mean daily maximum temperatures (mean T_max_) during early development was associated with reduced survival probabilities of young in all three development stages. No young survived when mean T_max_ > 38°C across all group sizes. Low reproductive success at high temperatures has broad implications for recruitment and population persistence in avian communities given the rapid pace of advancing climate change. That impacts of high temperatures were not moderated by group size, a somewhat unexpected result given prevailing theories around the influence of environmental uncertainty on the evolution of cooperation, suggests that cooperative breeding strategies are unlikely to be advantageous in the face of rapid anthropogenic climate change.

## Introduction

Anthropogenic climate change has altered weather patterns in every ecosystem on Earth [1,2], with far-reaching consequences for population dynamics across taxa [3]. An improved understanding of life history responses to current environmental variability is required to predict species-specific responses to climate change [4]. While cooperative breeders occur in diverse habitats [5,6], comparative research has demonstrated that both cooperatively-breeding birds [7] and mammals [8] occur with disproportionate frequency in regions characterised by high spatial and temporal variability in environmental conditions. This implies that group living enhances the ability to persist in challenging environments [9]. To date, however, there are few empirical studies that explicitly test the extent to which group-living mitigates the effects of climate variability on reproduction [10–13]. Of these studies, only Langmore *et al*. (2016) and van de Ven *et al*. (2019) explore impacts of temperature alongside variation in group size, despite evidence of thermoregulatory benefits of group living [14], and only Covas *et al*. [13] consider offspring survival across more than one development stage. This latter point is important, because specific drivers of survival can differ substantially between development stages [15–17].

Temperature and rainfall patterns are important measures of climate variability, playing a critical role in reproductive success rates observed in vertebrates [18], and recent changes in temperature and rainfall patterns have led to adjustments to the timing and success of reproduction in some bird species [19,20]. For birds in arid environments, higher rainfall is often associated with improved reproductive success [21,22], droughts are associated with reduced reproductive success [23,24] and periods of very hot weather are typically associated with lower nest survival rates [15,25] and nestling growth rates [26–28]. Therefore, it is reasonable to expect that reproductive success and population persistence of birds in arid environments will be impacted as regions become hotter and drier under climate change [29,30].

Cooperative breeding, where more than two individuals rear a single brood [31], occurs in ∼9% of bird species [32]. Reproductive benefits of cooperation include earlier fledging age and more broods raised per season [33]; reduced costs of breeding for females [11,34]; enhanced egg investment [35]; increased fledgling recruitment [17,36]; and the ability to raise overlapping broods [12,37]. Global comparative studies suggest that cooperative breeding evolved in unpredictable environments [38], facilitated the colonisation of such environments [39], or prevented extinction under increasingly harsh conditions [40]. One prominent explanation for the occurrence of cooperative breeding in birds in variable environments is that it represents a ‘bet-hedging’ strategy [38], whereby breeding individuals share the costs of reproduction with helpers and are thus able to reduce interannual variation in reproductive success in response to unpredictable rainfall and food availability [9]. This implies that cooperation might buffer breeding attempts from failure during adverse environmental conditions [13,41].

A likely mechanism underlying such benefits of cooperation is load-lightening [42], which refers to individual reductions in workload in response to the presence of additional group members. Load-lightening has been observed in a number of cooperatively-breeding species [11,16,43], and may operate via task-partitioning [37,44] or by improved access to resources [45,46]. In larger groups, there are more individuals available to assist with breeding attempts, which can either lead to load-lightening amongst individual group members [11,37], or to cumulatively greater investment in young [36,47]; both are potential benefits of group living that may be particularly advantageous when unfavourable rainfall or temperature conditions are experienced. Specifically, these effects could mean that larger groups are better able to maintain adequate levels of parental care to eggs, nestlings, and/or fledglings at high temperatures or during periods of low rainfall, despite individual declines in investment in parental care behaviours.

We use a comprehensive 15-year dataset on southern pied babblers *Turdoides bicolor* (hereafter ‘pied babblers’), a cooperatively-breeding passerine endemic to the Kalahari in southern Africa, to explore the impacts of temperature, rainfall, and adult group size (where group size indicated the number of potential adults available to help the breeding pair) on reproductive success, including fledgling survival. Specifically, we test for effects of these parameters on survival of young from 1) initiation of incubation to hatching; 2) hatching to fledging; and 3) fledging to nutritional independence at 90 days of age [48]. We expected high temperatures to reduce survival, and high rainfall and larger group sizes to enhance survival during each development stage. If the presence of helpers buffers the effect of environmental variation on reproduction, as proposed by the temporal variability hypothesis [9], then we would expect an interaction between environmental factors and group size, such that weaker impacts of adverse climatic conditions on reproduction are observed in larger groups.

## Materials and methods

### Study site and system

Fieldwork was conducted at the Kuruman River Reserve (33 km^2^, KRR; 26°58’S, 21°49’E) in the southern Kalahari. Mean summer daily maximum temperatures at the study site, from 1995-2015, averaged 34.7 ± 9.7°C and mean annual precipitation averaged 186.2 ± 87.5mm [49]. The Kalahari region is characterised by hot summers and periodic droughts [50], with extremely variable rainfall between years [51] and increases in both the frequency and severity of high temperature extremes over the last 20 years [52]. Pied babblers are medium-sized (60–90 g), cooperatively-breeding passerines endemic to the Kalahari where they live in territorial groups ranging in size from 3–15 adults [53]. They breed during the austral summer, from September to March [54]. Pied babbler groups consist of a single breeding pair with subordinate helpers [55], and all adult group members (individuals > 1 year old) engage in cooperative behaviours, including territory defence and parental care [48,54]. Previous research has shown that high temperatures and drought negatively affect many aspects of this species’ ecology, including foraging efficiency, body mass maintenance, and provisioning of young [56– 58].

Birds in the study population are marked as nestlings with a unique combination of metal and colour rings for individual identification, and are habituated to observation at distances of 1– 5 m [48]. Habituated groups are visited weekly during the breeding season to check group composition and record life history events, including breeding activity.

### Data collection

Data were collected for each austral summer breeding season from September 2005–February 2019 (14 breeding seasons in total).

#### Nest life history data

Nest monitoring (location of nests, determination of incubation, hatch, and fledge or failure dates, records of group size and brood size) followed Ridley & van den Heuvel [33]. Nests were located by observing nest-building, and incubation start, hatch and fledge dates were determined by checking nests every two to three days. Breeding attempts were considered to have failed when nests were no longer attended, or when dependent fledglings were not seen on two consecutive visits. Failure dates were calculated as the midpoint between the date of the last pre-fail nest/group check and the date when the nest was no longer attended or the fledgling was missing. In most cases, it was not possible to determine the proximate cause of nest failure or death, although common causes of nest failure in this species include predation, abandonment, and nestling starvation [53,59].

Group size (number of adults present in the group; range: 2–10, mean = 4.2 ± 1.5) was recorded for each nest incubated. Brood size was recorded 11 days after hatching (range: 1–5 nestlings, mean = 2.7 ± 0.8), when nestlings were ringed. We defined early development as the period between initiation of incubation and nutritional independence at 90 days of age [48]. Average time from initiation of incubation to hatching is 14 ± 1.2 days. Average time between hatching and fledging is 15.4 ± 1.7 days. Pied babbler are nutrionally independent (receiving < 1 feed per hour) by 90 days of age [48].

#### Sexing & nestling mass

Pied babblers are sexually monomorphic (Ridley, 2016) and molecular sexing was used to determine the sex of individuals (*sensu* Fridolfsson & Ellegren 1999). Blood samples were collected by brachial venipuncture and stored in Longmire’s lysis buffer. Nestlings were ringed, blood sampled, and weighed to 0.1 g on a top-pan scale 11 days post-hatching (Mass_11_).

#### Temperature and rainfall

Daily maximum temperature (°C) and rainfall (mm) data were collected from an on-site weather station (Vantage Pro2, Davis Instruments, Hayward, USA). Missing weather data from 2009, 2010, and 2011 were sourced from a nearby South African Weather Services weather station (Van Zylsrus, 28 km away), which produces significantly repeatable temperature measurements (Lin’s concordance correlation coefficient *r*_*c*_ = 0.957, 95 % CI: 0.951–0.962), and moderately repeatable rainfall measurements (*r*_*c*_ = 0.517, 95 % CI: 0.465–0.566) in comparison with the on-site weather station. Absolute differences in measured rainfall were small (average difference = 0.045 ± 3.075 mm, 95 % CI = −5.981–6.072 mm), suggesting that both weather stations adequately detected wet vs. dry periods.

Daily minimum (T_min_) and maxium (T_max_) temperatures, daily temperature variation (T_max_ −T_min_), were averaged for each development stage: incubation (mean T_minInc_, mean T_maxInc_, mean T_varInc_), nestling (mean T_minBrood_, mean T_maxBrood_, mean T_varBrood_), and fledgling (mean T_min90,_ mean T_max90,_ mean T_var90_). Rainfall was summed for the 60 days prior to initiation of incubation (Rain_60_), and for the period between fledging and independence (Rain_90_).

### Statistical analyses

Statistical analyses were conducted in R v 3.6.0 [62]. All continuous explanatory variables were scaled by centering and standardising by the mean [63,64]. All explanatory variables were tested for correlation with one another [65]. Mean T_max90_ and mean T_min90_ were correlated (VIF = 2.8) and all other explanatory variables were not correlated with each other (all VIF < 2). Correlated variables were not included in same additive models. Sample sizes reflect data sets after removing records containing missing values. Unless otherwise indicated, summary statistics are presented as mean ± one standard deviation. Analyses exclude groups > 8 due to small sample sizes for groups of 9 (*n* = 5) and 10 (*n* = 1) over 15 years of records. A quadratic term for temperature was included as a predictor variable only when no main linear effect of temperature was found and visual inspection of the data suggested a non-linear relationship. Sensitivity power analyses, using the package *pwr* [66], indicated that we had sufficient statistical power to detect effects of two-way interactions given our sample sizes [67,68], see Table S1. We tested for temporal trends in environmental (temperature and rainfall) and reproductive (nest success, fledgling survival) parameters using univariate linear models with breeding season as the only predictor. Covariates exhibiting temporal trends were detrended prior to inclusion in our models using the *detrend* function in the package *pracma* [69].

#### Survival probabilities during each development stage

Pied babbler survival probabilities are not constant across time during early development [54], see Fig. S1, and covariates are unlikely to have the same relationship with survival during all three early development stages. We used generalised linear mixed-effects models (GLMMs) with a binomial distribution and a logit link function in the package *lme4* [70] to determine which variables best predicted survival probabilities during each development stage. These analyses were undertaken at the level of the breeding attempt (i.e. clutch or brood) because individual offspring were only ringed for individual identification from the 11^th^ day after hatching, by which time ∼60% of monitored breeding attempts had failed. Model selection using Akaike’s information criterion corrected for small sample size (AICc), with maximum likelihood estimation was used to test a series of models to determine which model/s best explained patterns of variation in the data [63]. Where several models were within 5 AICc of the top model, top model sets were averaged using the package *MuMin* [71]. Model terms with confidence intervals not intersecting zero were considered to explain significant patterns in our data (Grueber, Nakagawa, Laws, & Jamieson, 2011). Binomial model fits were tested against the dispersion parameter in the package *RVAideMemoire* [73].

We considered the influence of the following parameters on (a) the probability of at least one egg per clutch surviving to hatch, (b) the probability of at least one nestling per brood surviving to fledge, and (c) the probability of at least one fledgling per brood surviving to nutritional independence: for (a) group size, Rain_60_, mean T_minInc_, mean T_maxInc_, and mean T_varInc_, for (b) group size, Rain_60_, mean T_minBrood_, mean T_varBrood_, mean T_maxBrood_, and mean T_maxBrood_^2^, and for (c) group size, Rain_90_, mean T_min90,_ mean T_var90,_ mean T_max90,_ and mean T_max90_^2^ We tested two-way interactions between rainfall, T_max_, and group size variables in each analysis, and, in order to account for non-independence of data, we included group identity as a random term in all three analyses.

In order to contribute data that can be incorporated into mechanistic modelling of the effects of climate change on avian populations [4], we further sought to identify the threshold T_max_ (‘breakpoints’) above which survival was compromised during each development stage. We used the package *segmented* [74] to apply a Davies’ test for a non-zero difference-in-slope and, when the regression parameter in the linear predictor was non-constant, to identify a point estimate and 95% confidence interval for the breakpoint. We then fitted simple linear regression models for the data above and below the identified breakpoints with temperature (mean T_maxInc,_ mean T_maxBrood,_ and mean T_max90_ respectively) as the ony predictor, and survival per breeding attempt as the response. In the segmented regressions, we used a continuous form of the survival response, specifically (a) the number of days between initiation of incubation and either the hatching of at least one egg or failure of the breeding attempt before hatching (age at hatch/fail), (b) the number of days between hatching and fledging at least one nestling from a brood or failure (age at fledge/fail), and (c) the number of days between fledging and at least one fledgling surviving to nutritional independence or failure (age at survival/fail).

#### Influence of nestling mass on fledgling survival

In addition to survival data at the scale of the breeding attempt, we have detailed individual-level survival data for 372 fledglings weighed and banded as 11-day-old nestlings. Larger nestling mass is commonly associated with higher survival probabilities in birds [16,75]. Prior research on pied babblers has shown that nestling mass is influenced by environmental factors such as temperature and rainfall [56]. We therefore used a confirmatory path analysis [76,77] to test for indirect effects of environmental and group size factors on survival to nutritional independence in known individual fledglings mediated via their mass as a nestling (Mass_11_). We computed the path analysis using the R package *piecewiseSEM* [78], which can accommodate multiple error structures. This capacity is important because the response terms of our component models have different distributions (see below). While model selection processes can be applied to multiple path analyses [79], our goal was not to choose between competing hypotheses, and but rather to construct a single model testing the relative importance of direct effects of environmental and group size factors vs. effects mediated via nestling mass. Path analysis allowed us to specify and simultaneously quantify all hypothesised relationships of interest, including the indirect effects of weather and group size on survival via nestling mass. Path coefficients are partial regression coefficients and can be interpreted similarly to simple and multiple regression outputs. Statistical significance was taken as p < 0.05. We hypothesised that:

Survival would be negatively affected by a) high temperatures during the nestling and fledgling stages, b) low rainfall between fledging and independence, c) smaller group size, and d) low nestling body mass (model with binomial error structure).

Nestling body mass would be negatively affected by a) high mean temperatures during the nestling period, b) low rainfall prior to the nestling period, and c) smaller group size (model with Gaussian error structure).

## Results

### Temporal patterns in temperature, rainfall and reproductive success

The total number of days (Oct–Mar) exceeding 35.5°C, identified as a critical temperature threshold in pied babblers [56,57], has increased significantly at the study site since 2005 (*F*_*1,12*_ = 7.448, *p* = 0.018; Fig.1a). Total summer rainfall (Oct–Mar) over the same time period was highly variable but showed a declining, non-statistically significant trend (*F*_*1,12*_ = 1.616, *p* = 0.228; Fig.1b). Both the number of nests fledged (a non-significant trend; *F*_*1,12*_ = 3.747, *p* = 0.077; Fig.1c) and the number of surviving young produced (*F*_*1,12*_ = 5,285, *p* = 0.040; Fig.1d) have declined at the study site since 2005, despite the number of groups monitored remaining relatively constant between years (coefficient of variation = 0.17). Most rain falls between Dec and Feb (72%), when temperatures are high (Fig.1e). Most pied babbler breeding activity occurs between Oct and Dec (68%), when conditions are generally drier and cooler than later in the season (Fig.1e).

**Figure 1:**
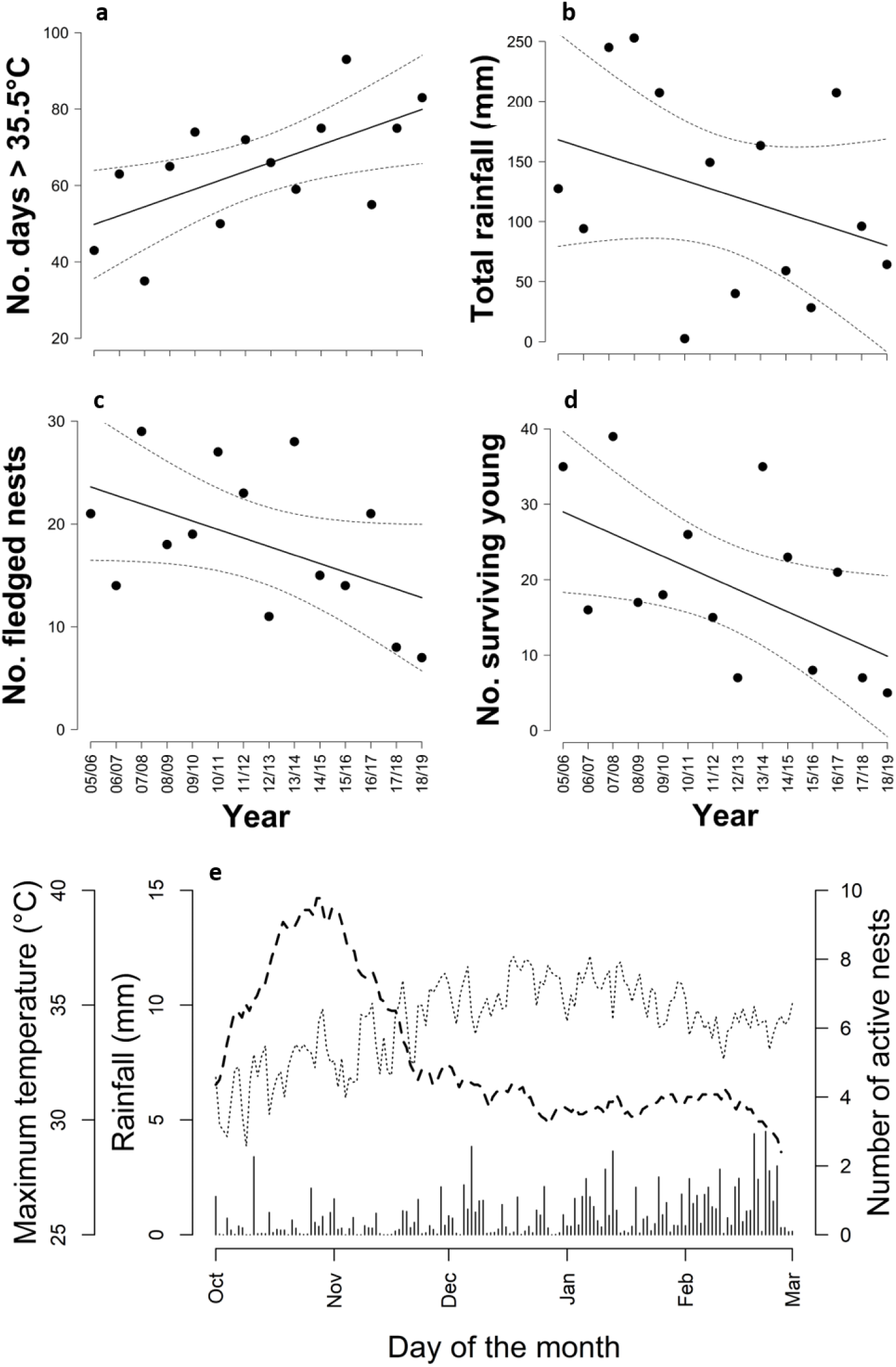
(a) the number of days > 35.5°C at the study site (b) total summer rainfall, (c) number of southern pied babbler *Turdoides bicolor* nests fledged in the study population, and (d) number of surviving young produced in the study population per breeding season per year (austral summer: 1 Oct to 1 Mar) since 2005. Black lines represent predictions from the models, and dashed lines the 95% CIs. (e) breeding activity between Oct and Mar (average number of active nests per day: dashed line), relative to temperature (average daily maximum temperature (°C) per day: dotted line) and rainfall (average rainfall (mm) per day: vertical bars).

#### Survival probabilities during each development stage

Overall, 31.4 ± 10.9% of breeding attempts produced at least one fledgling that survived to nutritional independence. Mean (± se) survival probabilities of young differed between life stages (incubation, nestling, dependent fledgling), with lower survival probabilities during early development in the nest, on average, than post-fledging (Fig. S1c).

Of 489 breeding attempts by 50 groups over 14 breeding seasons, 339 hatched (69.3%). The probability of at least one egg per clutch hatching decreased as mean daily maximum temperatures (mean T_maxInc_) increased (Table 1a). We found no evidence that mean T_minInc_, mean T_varInc_, rainfall or group size, or interactions between environmental conditions, influenced the probability of hatching (see Table S2 for full model output). For the period between initiation of incubation and hatching, a breakpoint was detected at 35.4°C (95% CI: 33.9, 36.9): there was no effect of mean T_maxInc_ on age at hatch/fail below 35.4°C (*F*_*1,399*_ = 0.008, *p* = 0.926), whereas above 35.4°C, age at hatch/fail significantly declined with increasing temperature (*F*_*1,85*_ = 9.490, *p* = 0.003, Fig. 2a).

**Table 1:**
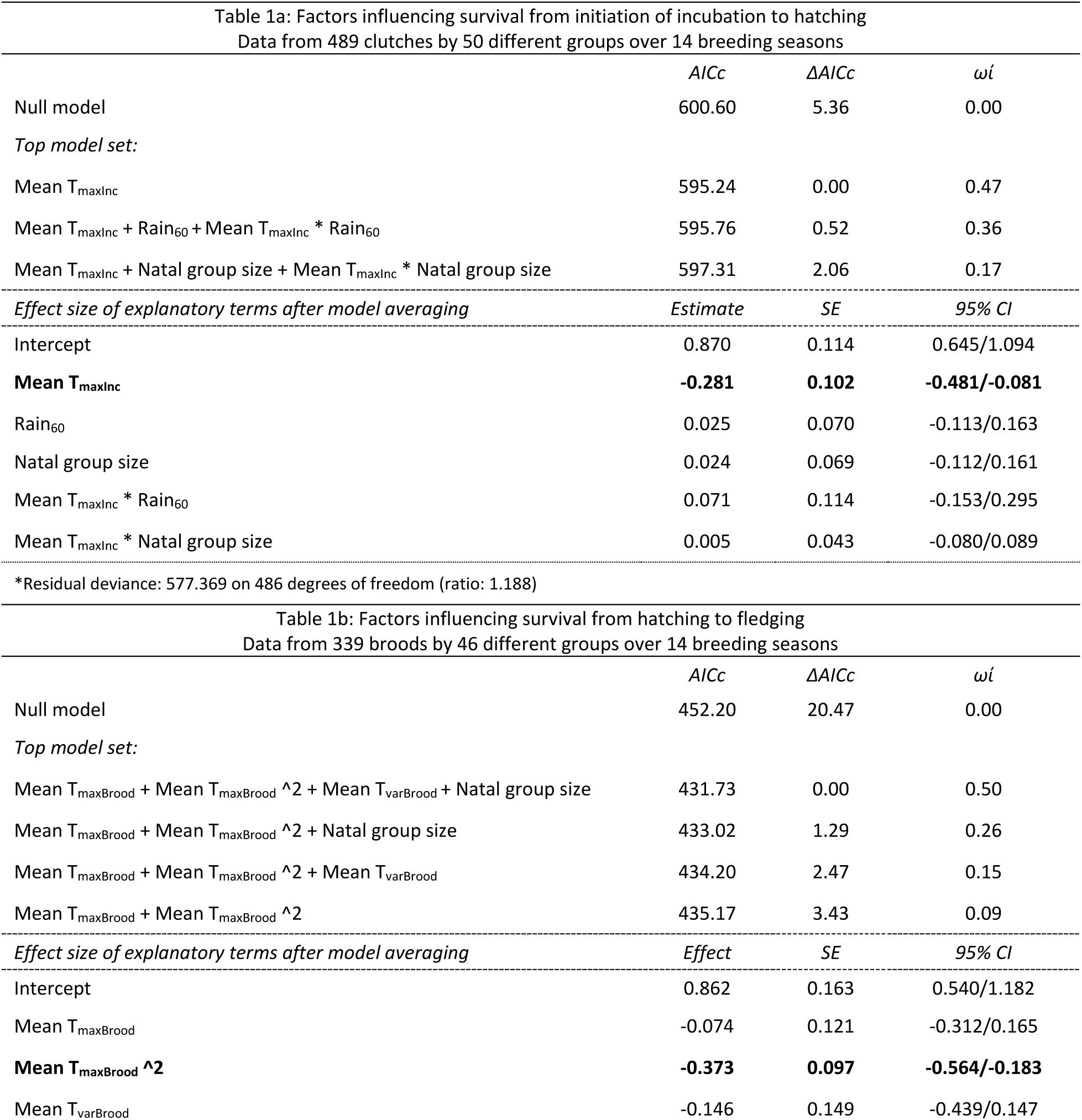
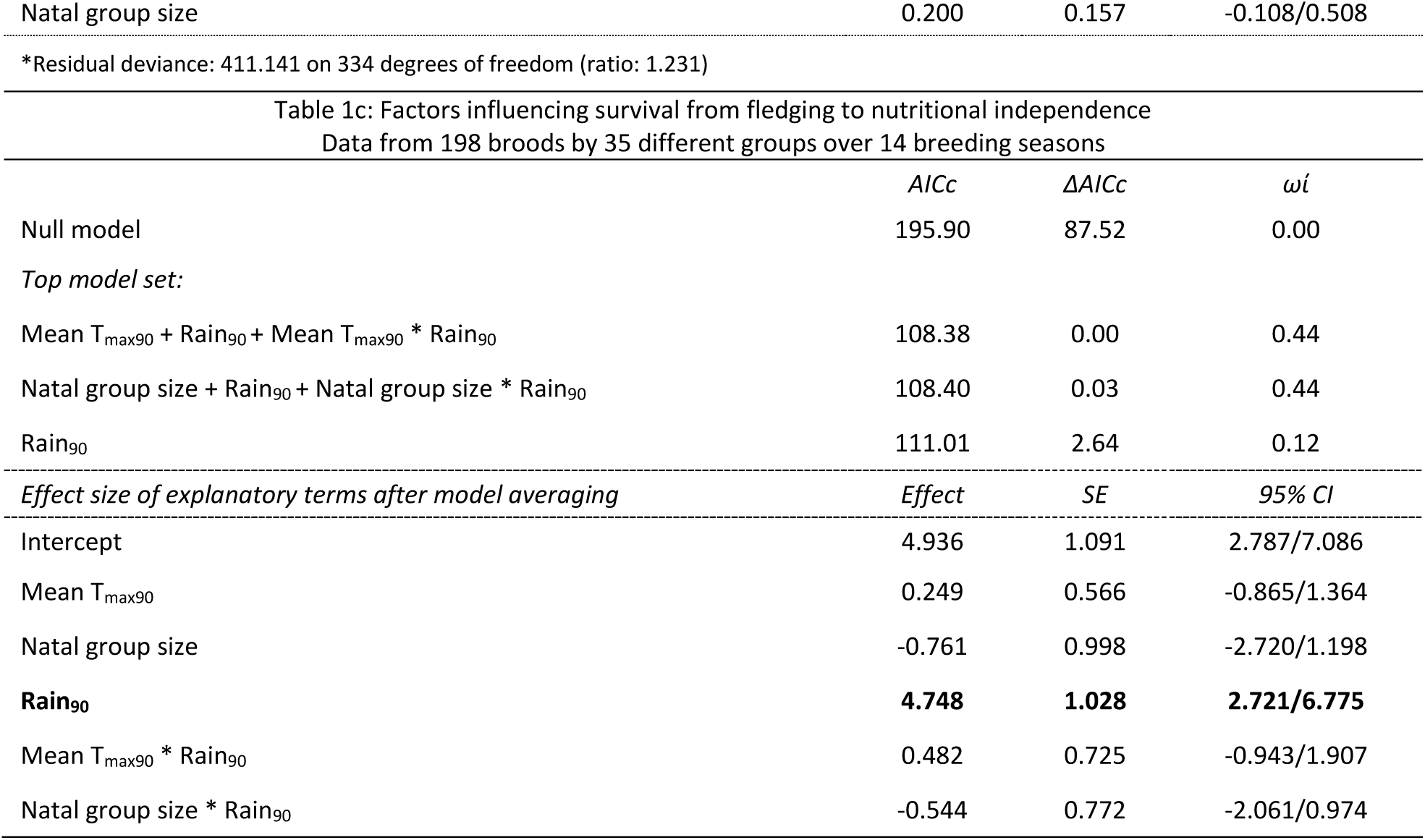
Top GLMM model set for factors influencing survival during early development. Model averaging was implemented for models with ΔAICc < 5 of the ‘best-fit’ model. Significant terms after model averaging are shown in bold. Null models shown for comparison with top model sets.

**Figure 2:**
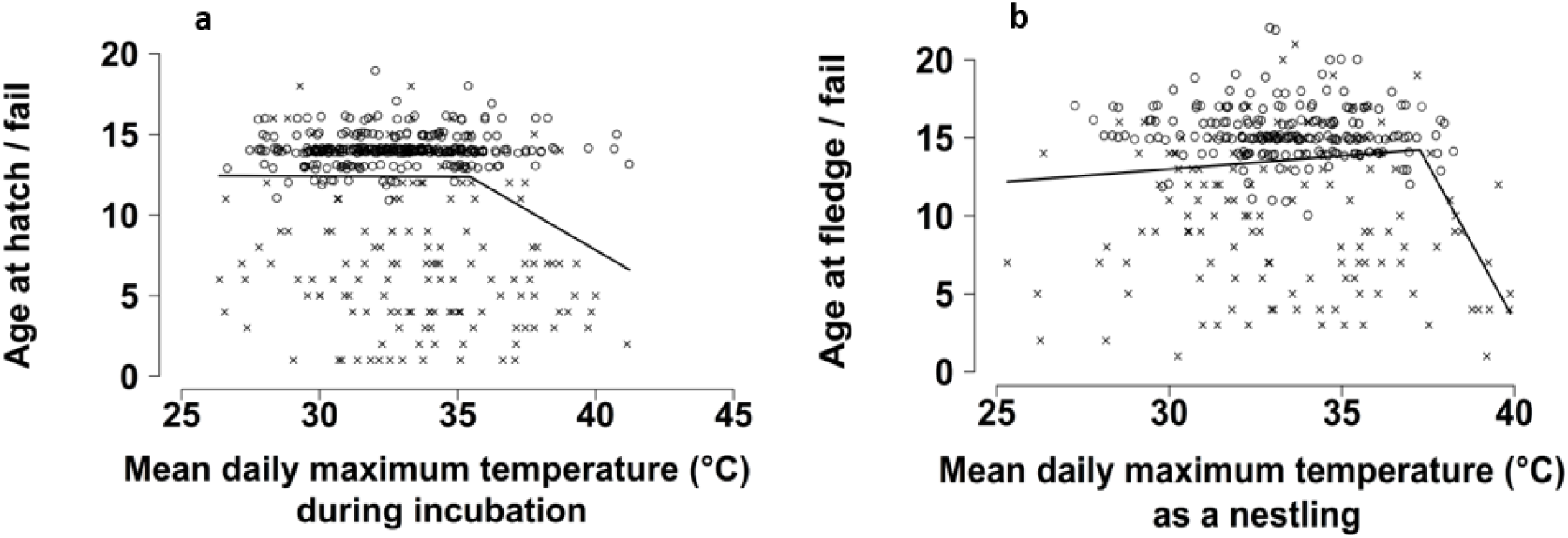
Survival from (a) initiation of incubation to hatch and (b) hatch to fledge as a function of mean daily maximum air temperature during the corresponding time period. Lines represent segmented linear regressions for the relationship between survival age and air temperature above and below the detected temperature thresholds. Open circles indicate that the clutch (a) or brood (b) transitioned to the next development stage; crosses indicate failure of the clutch (a) or brood (b).

Of 339 hatched nests by 46 groups over 14 breeding seasons, 210 fledged at least one chick (61.9%). The probability of at least one nestling per brood surviving to fledge increased with increasing mean daily maximum temperatures during the nestling period (mean T_maxBrood_) until ∼33.1°C, above which survival probability decreased (Table 1b). We found no evidence that mean T_minBrood_, mean T_varBrood_, rainfall or group size, or interactions between group size and environmental conditions, influenced the probability of fledging (see Table S3 for full model output). For the period between hatching and fledging a breakpoint was detected at 37.3°C (95% CI: 36.5, 38.0). Age at fledge/fail tended to increase with increasing mean T_maxBrood_ until 37.3°C (*F1,317* = 3.239, *p* = 0.073), above which age at fledge/fail declined significantly with increasing temperature (*F*_*1,20*_ = 13.370, *p* = 0.002, Fig. 2b). At mean T_maxBrood_ > 38°C (n=12), no nests fledged young.

Of 198 fledged broods with complete weather data by 36 groups over 14 breeding seasons, 160 produced at least one fledgling that survived to nutritional independence (80.8%). The probability of surviving to nutritional independence increased as rainfall during the post-fledging period increased (Rain_90_; Table 1c). We found no evidence that mean T_min90c_, mean T_max90_, mean T_var90_, group size or interactions between group size and environmental conditions influenced the probability of fledgling survival to independence (see Table S4 for full model output). We also found no evidence for a breakpoint in the data related to variation in mean T_max90_ for the period between fledging and independence (Davies’ test *p* = 0.288). While temperature was not a significant predictor of survival to nutritional independence in either of the model sets, no breeding attempts produced surviving young at mean T_max90_ > 38°C (n=8).

#### Influence of nestling mass on fledgling survival

The confirmatory path analysis model explained 47% of the variation in survival from fledging to independence (Fig. 3; *X*^2^ = 0.689, *p* = 0.708). Higher Rain_90_ was directly associated with an increased probability of surviving to independence, and larger group sizes were indirectly associated with increased survival via the positive effect of larger group size on nestling Mass_11_. High mean T_maxBrood_ was associated with reduced survival both directly and indirectly (high mean T_maxBrood_ was associated with reduced nestling mass, which in turn predicted reduced survival). There was no evidence for a direct effect of either mean T_max90_ or natal group size on survival to independence, or an effect of rainfall prior to the breeding attempt (Rain_60_) on nestling Mass_11_.

**Figure 3:**
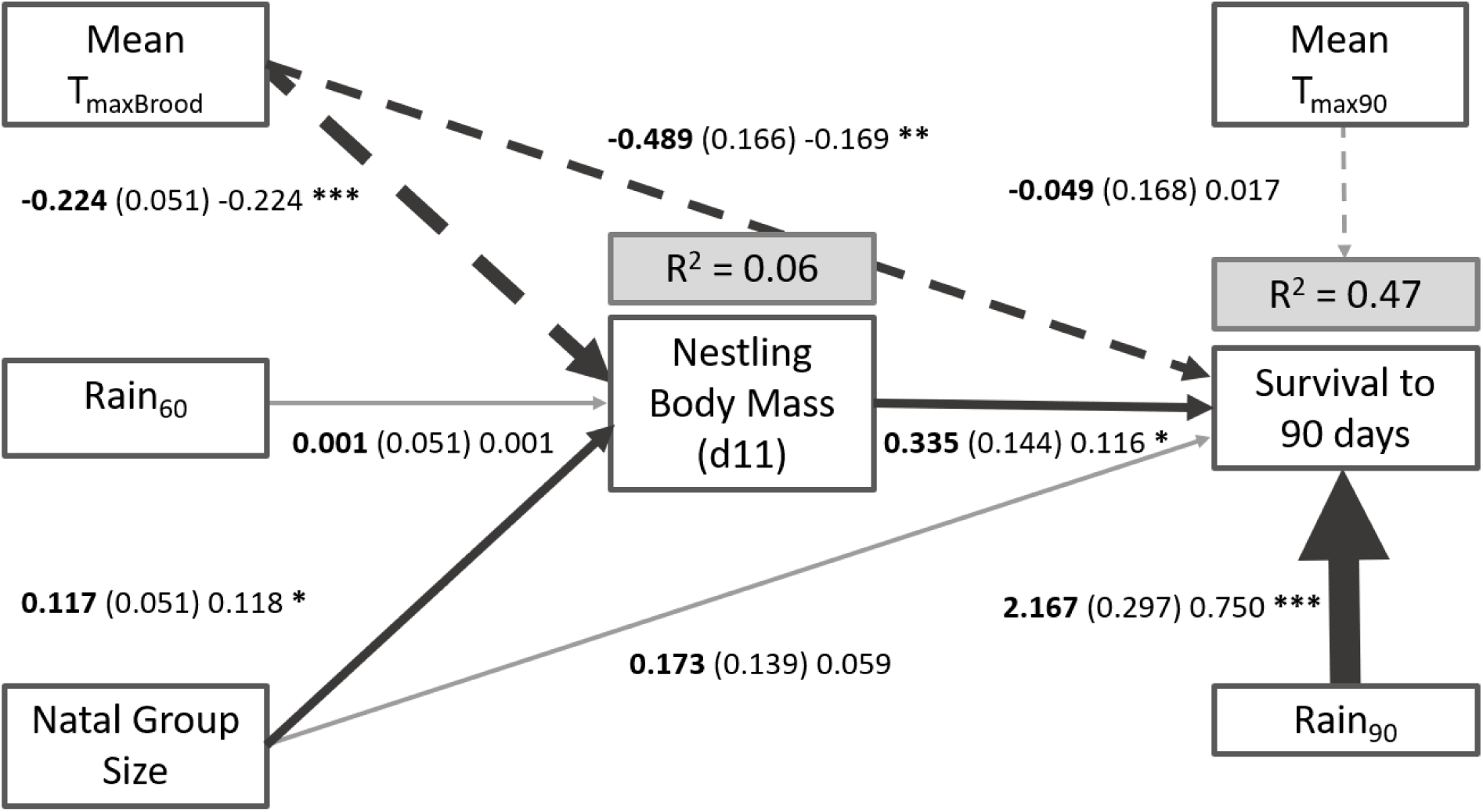
Confirmatory path analysis exploring the effects of environmental factors (temperature and rainfall) and group size on nestling body mass and survival to nutritional independence (90 days). Boxes represent measured variables. Arrows represent hypothesised unidirectional relationships among variables. Solid arrows denote positive relationships, dashed arrows negative relationships. Unstandardised path coefficients are shown in bold, followed by standard errors in parentheses, standardised estimates, and an indicator of statistical significance of the effect (*). Non-significant paths are grey. The thickness of significant paths has been scaled relative to the absolute magnitude of the standardised estimates, such that stronger effects have thicker arrows. R^2^ for component models are given in the grey boxes above response variables.

Specifically, larger nestlings were more likely to survive to independence (Est = 0.116, *p* = 0.019), as were fledglings that experienced higher Rain_90_ (Est = 0.750, *p* < 0.001). However, fledglings were less likely to survive (Est = −0.169, *p* = 0.003) when they had experienced higher mean T_maxBrood_. Nestlings were heavier when raised by larger groups (Est = 0.118, *p* = 0.021) and lighter when they experienced higher mean T_maxBrood_ (Est = −0.224, *p* < 0.001). There was an indirect negative effect of mean T_maxBrood_ on survival via nestling mass [Est = −0.026 (calculated by multiplying standarised estimates for each component of the indirect path: −0.244 × 0.116; see Fig. 3)]. The combined direct and indirect effect of mean T_maxBrood_ (via nestling mass) on survival was negative [-0.195, calculated by summing standardised estimates for direct and indirect paths: −0.169 + (−0.026); see Fig. 3]. The direct effect of mean T_maxBrood_ was more prominent (∼87% of the combined effect) than the indirect effect via nestling mass (∼13% of the combined effect). Natal group size had an indirect positive effect on survival via nestling mass [Est = 0.014 (= 0.118 × 0.116)], with an overall effect of natal group size = 0.132 (0.059+0.014).

## Discussion

We investigated the impacts of environmental conditions and the potential for group living to buffer against these impacts in a cooperatively breeding bird, focusing on egg, nestling, and fledgling survival. We present three main findings. First, exposure to high mean daily maximum temperatures during early development was associated with significant reductions in survival probabilities, in keeping with other recent studies [24,26,80]. Second, both environmental (largely direct) and social (indirect) factors were important for predicting survival during different development stages. Third, contrary to our expectations, we found no evidence that effects of T_max_ and rainfall on reproduction were moderated by group size, despite considerable statistical power to detect such interactions. Taken together with evidence that temperatures are increasing and rainfall decreasing at the study site, and that the number of surviving young produced in the study population showed similar declines, the impacts of high temperatures on mortality during early development are concerning.

### Impacts of high temperatures during early development

In pied babblers, high mean daily maximum temperatures during early development were associated with a significantly increased risk of mortality. Adverse weather is known to impair egg [81] and nestling development [82]. For example, survival to fledging can be compromised both by sub-optimally cool [83] and sub-optimally hot conditions [26,28]. For the early development stages before fledging, we identified temperature thresholds in the mid-to high-30s (35.4°C during incubation, and 37.3°C for nestlings) above which survival of eggs and young became significantly less likely. These temperatures are within ∼2°C of an apparent upper limit (38°C) above which we recorded no successful breeding in this species over 15 years of research. While we did not detect a direct effect of high temperatures on post-fledging mortality (i.e. between fledging and independence) at the brood scale, path analysis revealed that the probability of individual fledglings surviving to independence was influenced by high temperatures experienced as a nestling (Mean T_maxBrood_), both directly and indirectly via the effect of high temperatures on nestling Mass_11_. This suggests that dependent pied babbler fledglings, similar to the young of other species [84,85], are influenced by carryover effects of high temperatures they experienced while still in the nest. With temperatures increasing rapidly in the Kalahari (van Wilgen *et al*. 2016), the 38°C limit for successful breeding in this species suggests that pied babblers may increasingly experience conditions that do not allow successful breeding. This could undermine population growth and ultimately lead to local extinctions for this species.

### Different drivers of survival for each early development stage

The primary climatic (temperature and rainfall) and social (group size) drivers of survival probability were different across the three development stages. Mean daily maximum temperature was the strongest predictor of survival probability during both the incubation and nestling development stages. At high temperatures over prolonged periods, incubating birds may not be able to sustain nest attendance to regulate egg temperature [86], leaving eggs vulnerable to overheating and becoming unviable [15,81]. Likewise, several studies have reported that high temperatures constrain nestling growth [26,82,87], result in smaller nestlings overall [27,88,89], alter corticosterone levels [90,91], and reduce nestling survival probabilities [92,93].

Rainfall was the strongest predictor of survival probability during the dependent fledgling stage. Higher rainfall periods are associated with greater food availability [21,94], which likely enhanced both provisioning rates to fledglings [95] and their ability to find food for themselves [96,97]. In cooperative breeders, survival of young during this stage often improves with increasing group size [36,54]: larger groups may provision more regularly (Meade *et al*. 2010; but see Wiley & Ridley 2016), better detect and repel predators [53], or access higher quality territories or nest sites [16]. We did not find a direct effect of group size on survival to independence. However, path analysis indicated that group size influenced individual fledgling survival probabilities indirectly, via a positive effect on nestling mass - itself a well-established positive predictor of post-fledging survival in cooperative breeders [16,75]. The presence of both direct and indirect (via negative effects on nestling Mass_11_) effects of mean T_max_ during the nestling period on survival to independence suggests that carryover effects of high temperatures during early development continue to impact individual survival probabilities post-fledging [98,99].

### Buffering effect of group size

We found no evidence that group size mediated effects of high mean T_max_ on breeding outcomes. While we found an indirect positive effect of larger group size on survival from fledging to independence, group size did not interact with the large and persistent negative effects of high mean T_max_ on survival observed across all development stages to buffer the detrimental effects of high temperatures on survival from one early development stage to the next. This suggests that physiological tolerance limits [100] and resource constraints [101] at high temperatures may exceed any potential buffering effect of group size on reproductive success in cooperative breeders in arid and semi-arid environments [10].

### Conclusion

In this study, negative effects of adverse climate conditions on breeding success in a cooperative breeder were not moderated by group size, suggesting that reproduction in pied babblers is constrained by available resources and physiology at high temperatures and low rainfall, regardless of group size. Climate change is one of the defining challenges of our time, posing a serious threat to biodiversity [3] and society [1]. The Intergovernmental Panel on Climate Change now predicts with virtual certainty that the incidence of hot extremes will continue to become more frequent and the length, frequency, and intensity of heatwaves will continue to increase over most land masses [102]. At higher average and extreme temperatures, arid zone bird species may increasingly experience temperatures that preclude successful breeding. We have observed both increasing temperatures and declining rainfall, along with declining reproductive success at the study site over the last 15 years. Over time, the negative effects of these high temperatures on offspring survival could limit population recruitment and lead to local extinctions. Despite the intuitive appeal of the hypothesis that cooperative breeding should buffer against some of these effects, we found no evidence this will be the case in pied babblers. Taken together, our findings raise concerns for the long-term persistence of arid zone species in the face of rapidly changing environmental conditions, and suggest that cooperative breeding strategies are unlikely to confer an advantage over alternative breeding strategies as species respond to advancing climate change.

## Acknowledgements

We thank the Kuruman River Reserve (KRR) and surrounding farms, Van Zylsrus, South Africa, for making the work possible. The KRR was financed by the Universities of Cambridge and Zurich, the MAVA Foundation, and the European Research Council (Grant No. 294494 to Tim Clutton-Brock), and received logistical support from the Mammal Research Institute, University of Pretoria. Thanks to Sello Matjee, Paige Ezzey, and Lesedi Moagi for fieldwork during 2016– 2019, and all past and present staff and students of the Pied Babbler Research Project for data collected since 2003. Work was funded by the DST-NRF Centre of Excellence at the FitzPatrick Institute of African Ornithology, the University of Cape Town, the Oppenheimer Memorial Trust (Grant No. 20747/01 to ARB), the British Ornithologists’ Union, the Australian Research Council (Grant No. FT110100188 to ARR), and the National Research Foundation of South Africa (Grant No. 99050 to SJC). The opinions, findings and conclusions are those of the authors alone, and the National Research Foundation accepts no liability in this regard. Data were collected under animal ethics permits R2012/2006/V15/AR and 2016/V6/SC, University of Cape Town. We thank the associate editor and two anonymous reviewers for their thorough and thoughtful comments which helped us to improve the manuscript immeasurably.

## Notes

### Competing Interest Statement

The authors have declared no competing interest.

