## Supplementary material for "High temperatures drive offspring mortality in a cooperatively breeding bird": Table S1

### Appendix 1: Supporting Information

### Power analyses for the interactions

Interactions between group size and climatic effects on development would be consistent with a buffering effect of group size on survival. We therefore conducted sensitivity power analyses to identify the minimum determinable effect of two-way interactions given our sample sizes (Cohen, 1988; Greenland et al., 2016), using the package *pwr* (Champely et al., 2018). For our regression models, we used the function **pwr.f2.test(u =,v =,f2 =,sig.level =,power =)**, where u = numerator degrees of freedom, v = denominator degrees of freedom, α (the signficance level representing the probability of finding an effect that is not there) = 0.05, and power (probability of finding an effect that is there) = 0.8. The value $f^2$ is the calculated value, representing the measure of determinable effect size. We assumed a fourfold increase in required sample size to adequately detect interactions in mixed-effects models (Leon & Heo, 2009), and confirmed we have sufficient sample size to detect a range of effect sizes, from small to large, in all analyses including main effects (all Cohen's $f^2 < 0.03$) and two-way interactions (all $f^2 < 0.12$) – see Table S1below. Cohen (1988) suggested that $f^2$ values of ~ 0.02, ~0.15, and ~0.35 represent small, medium, and large effect sizes respectively.

**Table S1**

***Power analyses for the interactions: multiple regression power calculations***

| Development stage | *u* | *v* | *α* | *power* | $f^2$ |
|---|---|---|---|---|---|
| **Egg** | | | | | |
| Main effects | 3 | 492 | 0.05 | 0.8 | 0.019 |
| Interactions | 3 | 123 | 0.05 | 0.8 | 0.080 |
| **Nestling** | | | | | |
| Main effects | 3 | 341 | 0.05 | 0.8 | 0.029 |
| Interactions | 3 | 85 | 0.05 | 0.8 | 0.118 |
| **Fledgling** | | | | | |
| Main effects | 3 | 378 | 0.05 | 0.8 | 0.026 |
| Interactions | 3 | 95 | 0.05 | 0.8 | 0.105 |

*u = model degrees of freedom; v = sample size, α = the significance level, and power (p) = probability of finding an effect that is there; $f^2$ = measure of determinable effect size (values of ~ 0.02, ~0.15, and ~0.35 represent small, moderate, and large determinable effect sizes respectively).

### Survival probabilities are not constant across time during early development

We used an exploratory Cox proportional hazards model (Cox, 1972) to visualise the relationship between overall risk of mortality (Austin, 2017) over time during early development and confirm the patterns identified by Ridley (2016) showing that survival probabilities are lower during the incubation and nestling period than after fledging. Setting survival probabilities per breeding attempt (i.e. per clutch or brood) as the response in the Cox model, we included the following parameters as predictor variables:

- pair tenure (the number of consecutive breeding seasons in which the same dominant breeding pair were present in a group),

- mean $T_{maxTotal}$ (the average daily maximum temperature for the whole dependent period from initiation of incubation until independence)
- $Rain_{Total}$ (sum of daily rainfall totals for the whole dependent period from initiation of incubation until independence)
- group size, and
- the interactions between i) mean $T_{maxTotal}$ and group size, ii) $Rain_{Total}$ and group size, and iii) mean $T_{maxTotal}$ and $Rain_{Total}$.

In order to account for non-independence of data, group identity and year were included as random effects.

A Cox proportional hazards analysis expresses mortality risk at each time step as a hazard ratio (HR), where HR > 1 indicates a higher risk of mortality. Model terms with HR confidence intervals not intersecting one are considered to explain significant patterns within the data. We computed the Cox model in the *survival* package (Therneau & Lumley, 2009), and visualised the model output using the *survminer* package (Kassambara, Kosinski, Biecek, & Scheipl, 2017).

The Cox proportional hazards model showed that overall mortality risk during the whole perid of early development was a) influenced by a combination of group size, temperature, and rainfall, and b) was not consistent for all development stages. Specifically, risk of mortality was higher for clutches and broods (i.e. while young were still in the nest) than for fledglings (Fig. S1a, n = 488 breeding attempts over 14 seasons). High temperatures during early development (HR = 1.145, 95% CI: 1.011,1.296, $z = 2.139$; including the quadratic term for temperature HR = 1.349, 95% CI: 1.275,1.426, $z = 10.472$; Fig. S1b) were associated with an increased risk of failure for breeding attempts (Fig. S1b). At mean daily temperatures ≥ 38°C ($n = 17$), 100% of breeding attempts failed. At the 17 coolest nests, where mean daily temperatures ≤ 28 °C, the proportion of nests that failed was also very high (~80%). This compares to an optimum temperature of ~34°C (*n* = 84) at which only 52% of breeding attempts failed. Breeding attempts undertaken by larger groups (HR = 0.699, 95% CI: 0.585,0.834, *z* = -3.980) and during wetter periods (HR = 0.297, 95% CI: 0.236,0.373, *z* = -10.372; Fig. S1b) were less likely to fail. When rainfall > 189 mm over the development period (*n* = 44), 0% of nests failed. Pair tenure (HR = 0.975, 95% CI: 0.859,1.107, *z* = -0.385) and the interactions between temperature and group size (HR = 0.975, 95% CI: 0.891,1.066, *z* = -0.562) and between rainfall and temperature (HR = 0.966, 95% CI: 0.827,1.129, *z* = -0.434) were not important for predicting risk of failure of breeding attempts (Fig. S1b). While an interaction between rainfall and group size was detected (HR = 0.782, 95% CI: 0.643,0.952, *z* = -2.453), further investigation indicated that this interaction was not robust (see below).

Overall, 31.4 ± 10.9% of breeding attempts produced at least one fledgling that survived to nutritional independence. Mean (± se) survival probabilities of young differed between life stages (egg, nestling, dependent fledgling; Fig. S1c). On average, survival during early development was lower (67.9 ± 10.4% of incubated clutches hatched; 62.2 ± 10.6% of hatched nests fledged) than after fledging (84.1 ±12.4% of broods produced at least one fledgling that survived one week; 88.3 ± 15.1% of broods that produced at least one week-old fledgling produced at least one independent juvenile).

From the Cox proportional hazard model, it was not possible to evaluate the relative influence of each significant predictor during each development stage, or to determine whether predictors influenced variation in mortality risk in the same way during each development stage. Therefore, in the main text of the manuscript, we only present the GLMM and confirmatory path analyses that directly address the influence of temperature, group size and rainfall during each development stage.

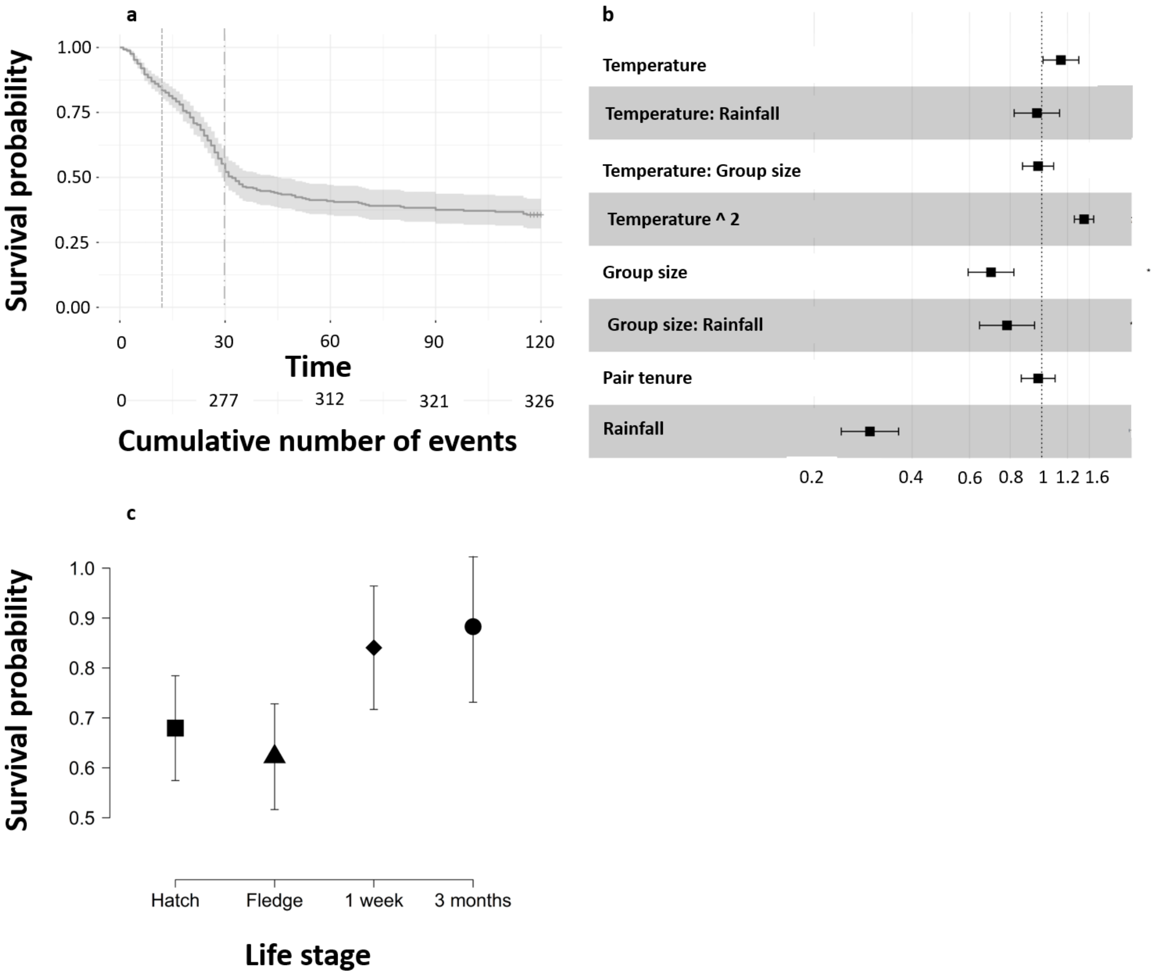

*Figure S1: (a) the estimated survival probability curve of young at the mean values of all covariates in the full Cox proportional hazards model between initiation of incubation and hatching (dotted vertical line) and between hatching and fledging (dashed vertical line; n = 488 nests). (b) forest plot showing the hazard ratio and 95% confidence interval for each covariate in the full Cox regression. Covariates with confidence intervals not crossing 1 are considered to explain significant patterns in the data. Hazard ratios above 1 indicate a positive association with probability of dying. (c) mean ± sd survival probabilities for each life stage transition: from start of incubation to hatch (square), from hatch to fledge (triangle), from fledge to 7 days of age (diamond), and from 1 week of age to nutritional independence at 3 months of age (circle).*

Although the Cox model identified a significant interaction between group size and rainfall, whereby larger groups required less rain in order to reproduce successfully than smaller groups (HR = 0.782, 95% CI: 0.643,0.952, $z$ = -2.453; Fig. S1b), visualisation of the data (Fig. S2) suggests the slopes are similar across group sizes. The observed relationship, that larger groups required less rain than smaller groups in order to breed successfully, is driven primarily by the higher survival of young in very large groups (8 individuals) at low rainfall, and their shallower increase in survival of young as rainfall increased, relative to all other group sizes. Groups with as many as eight adults are unusual and therefore seldom relevant in our study population. In this study, we observed only eight nests where group size = 8 (over 15 years of records). At group sizes < 8, group size did not appear to influence the shape of the relationship between rainfall and survival of young. We found no further evidence for an interaction between group size and rainfall in the finer-scale analyses for each early development stage. We therefore conclude that the observed interaction fails to provide strong evidence in support of a buffering effect of larger group size.

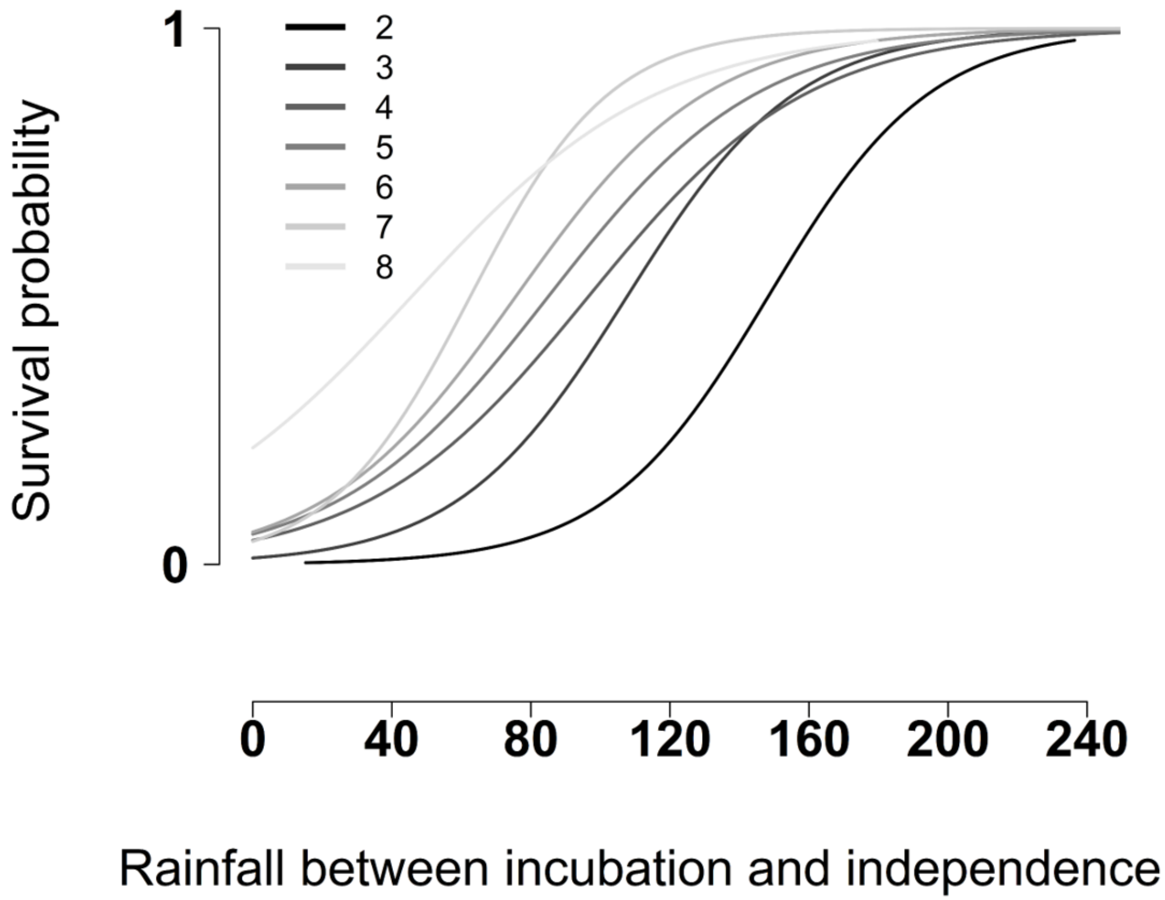

*Figure S2: There is a main effect of rainfall, with higher survival rates of breeding attempts at higher values of rainfall, and a main effect of group size, with higher survival rates of breeding attempts in larger groups, but the interaction effect detected in the Cox proportional hazards model is an artifact of the higher survival of young in very large groups (8 individuals) at low rainfall, and their shallower increase in survival of young as rainfall increased, relative to all other group sizes. Group sizes this large are seldom relevant in this study population (n = 8 over 15 years of monitoring).*

**Full GLMM model output tables for survival probability analyses for each development stage**

### Table S2

***Effect of group size and environmental factors on survival from initiation of incubation to hatching***

Data from 489 breeding attempts by 50 groups over 14 breeding seasons
Random terms: Group identity

| Model Term | AICc | ΔAICc | weight |
|---|---|---|---|
| Null model | 600.6 | 5.35 | 0.029 |
| Mean $T_{minInc}$ | 601.0 | 5.72 | 0.024 |
| Mean $T_{maxInc}$ | 595.2 | 0.00 | 0.146 |
| Mean $T_{varInc}$ | 601.1 | 5.86 | 0.022 |
| Natal group size | 601.0 | 5.73 | 0.024 |
| $Rain_{60}$ | 602.6 | 7.34 | 0.011 |
| Mean $T_{maxInc}$ * Natal group size | 597.3 | 2.06 | 0.148 |
| Mean $T_{maxInc}$ * $Rain_{60}$ | 595.8 | 0.52 | 0.320 |
| $Rain_{60}$ * Natal group size | 603.4 | 8.14 | 0.007 |

### Table S3

***Effect of group size and environmental factors on survival from hatching to fledging***

Data from 339 hatched nests by 46 groups over 14 breeding seasons
Random terms: Group identity

| Model Term | AICc | ΔAICc | weight |
|---|---|---|---|
| Null model | 452.2 | 20.49 | 0.000 |
| Mean $T_{minBrood}$ | 454.2 | 22.43 | 0.000 |
| Mean $T_{maxBrood}$ | 454.2 | 22.45 | 0.000 |
| Mean $T_{maxBrood}$ ^2 | 435.2 | 3.43 | 0.090 |
| Mean $T_{varBrood}$ | 447.5 | 5.81 | 0.000 |
| Natal group size | 449.0 | 17.22 | 0.000 |
| $Rain_{60}$ | 453.6 | 21.83 | 0.000 |
| Mean $T_{maxBrood}$ + Mean $T_{maxBrood}$ ^2 + Natal group size | 433.0 | 1.29 | 0.263 |
| Mean $T_{varBrood}$ + Natal group size | 443.9 | 12.15 | 0.001 |
| Mean $T_{maxBrood}$ + Mean $T_{maxBrood}$ ^2 + Mean $T_{varBrood}$ | 434.2 | 2.47 | 0.146 |
| Mean $T_{maxBrood}$ + Mean $T_{maxBrood}$ ^2 + Mean $T_{varBrood}$ + Natal group size | 431.7 | 0.00 | 0.500 |
| Mean $T_{maxBrood}$ * Natal group size | 451.3 | 19.52 | 0.000 |
| Mean $T_{maxBrood}$ * $Rain_{60}$ | 456.5 | 22.45 | 0.000 |
| $Rain_{60}$ * Natal group size | 452.3 | 20.59 | 0.000 |

**Table S4**

***Effect of group size and environmental factors on survival from fledging to nutritional independence***

Data from 198 broods from 35 groups over 14 breeding seasons
Random terms: Group identity

| Model Term | AICc | ΔAICc | weight |
|---|---|---|---|
| Null model | 195.9 | 87.54 | 0.000 |
| Mean $T_{min90}$ | 197.8 | 89.43 | 0.000 |
| Mean $T_{max90}$ | 197.6 | 89.27 | 0.000 |
| Mean $T_{var90}$ | 197.1 | 88.69 | 0.000 |
| Natal group size | 197.2 | 88.79 | 0.000 |
| $Rain_{90}$ | 111.0 | 2.64 | 0.119 |
| Mean $T_{max90}$ * Natal group size | 200.9 | 92.49 | 0.000 |
| Mean $T_{max90}$ * $Rain_{90}$ | 108.4 | 0.00 | 0.444 |
| $Rain_{90}$ * Natal group size | 108.4 | 0.03 | 0.437 |
